# *Smpd1* regulates chitin-clearance for tracheal gas-filling in the *Drosophila* embryo in a ceramide-specific manner

**DOI:** 10.1101/2024.04.18.589892

**Authors:** Alexander J Hull, Magda L Atilano, Jenny Hallqvist, Wendy Heywood, Kerri J Kinghorn

## Abstract

Type A and B Niemann Pick (NPD) is an inherited multisystem lysosomal storage disorder caused by mutations in the *SMPD1* gene. Respiratory dysfunction is a key hallmark of NPD, although the precise mechanisms underlying these pathologies is underexplored. Here we present a *Drosophila* model of *Smpd1* loss-of-function that displays significant respiratory defects. *Smpd1* is expressed in the late-embryonic fly respiratory network, the trachea, and is secreted into the tracheal lumen. Loss of *Smpd1* results in embryonic lethality, and although tracheal morphology appears normal, trachea fail to fill with gas prior to eclosion. We demonstrate that clearance of luminal constituents through endocytosis prior to gas-filling is defective in *Smpd1* mutants. This is coincident with autophagic, but not lysosomal defects. Finally, we show that although bulk sphingolipids are unchanged, dietary loss of lipids in combination with genetic and pharmacological block of ceramide synthesis is sufficient to rescue gas-filling defects. In summary, we present a novel NPD model amenable to genetic and pharmacological screens, and highlight myriocin, an inhibitor of ceramide synthesis, as a potential therapeutic drug for the treatment of NPD.

## Introduction

Type A and B Niemann Pick disease (NPD) is a multisystemic inherited disorder of metabolism caused by bi-allelic mutations in the *SMPD1* gene, of which more than 25 pathogenic variants have been identified (Zampieri et al., 2016, Stenson et al., 2020). *SMPD1* encodes the lysosomal hydrolase acid sphingomyelinase (ASM), which cleaves the sphingolipid sphingomyelin (SM) into ceramide and phosphocholine. In Type A and B NPD, SM accumulates in the lysosomes of engorged macrophages, termed foam cells, which infiltrate the reticuloendothelial system of various tissues, such as the liver, lungs, spleen, bone marrow and brain, causing multi-organ disease (McGovern et al., 2017). Moreover, heterozygous *SMPD1* mutations were recently discovered to be a potent Parkinson’s disease (PD) risk factor, although the exact mechanisms linking the two conditions are yet to be elucidated (Gan-or et al., 2013, 2015, Robak et al., 2017). As well as being linked via genetic studies, SM has been found to be enriched in Lewy bodies, intracytoplasmic neuronal inclusions of aggregated alpha-synuclein protein, the main neuropathological hallmark of PD (den Jage, 1969, Spillantini et al.,1998).

ASM functions at low pH principally within the lysosome, hydrolysing SM in a zinc-independent manner as part of the lipid salvage pathway. It is also trafficked through the secretory pathway to act extracellularly on SM at the plasma membrane in a zinc-dependent manner (Schissel et al., 1996). As expansion of the lysosomal compartment is the hallmark Niemann-Pick pathology, it is thought lysosomal ASM is the principal contributor to the disease, yet the contribution of secreted ASM to Niemann-Pick or PD pathology has yet to be effectively resolved (Kornhuber et al., 2015). Several studies have demonstrated altered plasma membrane composition in *SMPD1* mutant primary neurones, suggesting that functions at the membrane may be important to NPD progression (Galvan et al., 2008, Schuchman, 2010).

Although encouraging strides are being made for enzyme replacement therapy for Type A and B NPD with Olipudase Alfa, this therapy is still in its infancy and thus there is an urgent need to identify novel therapeutic targets that reverse lysosomal and downstream pathologies (Lachmann et al., 2023). Here we present a *Drosophila* model of *SMPD1* loss-of-function, mutated in the fly orthologue of *SMPD1, Smpd1*. We show that loss of ASM function leads to embryonic lethality and developmental defects in the respiratory system in *Drosophila*, mimicking those in patients with Type A and B NPD. We further demonstrate these defects can be rescued by dietary, genetic and pharmacological manipulations that decrease lipid and sphingolipid abundance.

## Materials and Methods

### Fly stocks and culture

Unless otherwise described, flies were raised on SYA food and maintained at 25°C and 50% relative humidity on a 12:12 light-dark cycle. For embryo collections, with the exception of dietary modifications, flies were placed onto grape juice agar plates, and following synchronisation, embryos were allowed to develop at 25°C (12hr for stage 16, 23hr for late stage 17). Holidic medium was prepared as previously described (Piper et al., 2014). For drug experiments, myriocin was dissolved in DMSO and added to the food at a final concentration of 100µM.

Unless otherwise stated, flies were backcrossed over 6 generations into a *w*^*1118*^ background to reduce background effects. The following stocks were obtained from Bloomington: *Smpd1*^*CRIMIC*^ (#91346), *Smpd1*^*KO*^ (#81092), *Smpd1*^*R571L*^ (#81093), *Lace*^*k5305*^ (#12176), *Lace*^*8*^ (#25150), *Schlank*^*G0061*^ (#11665), *Sply*^*05091*^ (#11393) *Lace*RNAi (#51475), Btl-GAL4 (#78328), Hemo-Gal4 (#78565), UAS-mCherry.nls (#38424), UAS-GFPmChAtg8a (#37749), UAS-preproANF::GFP (#7001), UAS-VermRFP (#86530), UAS-Cht-Tom (#66512), *Smpd1* RNAi 2 (#36760) and Srp-Hemo-moe.mCherry (#78362). The following stocks were taken from VDRC: sfGFP-CG3376 (#318772), SchlankRNAi (#v41114) and *Smpd1* RNAi 1(#v12227). *Smpd1*^*k*^ was from Kyoto stock centre (#104944).

Cpes^KO^ and UAS-Cpes were gifts from Jairaj Acharya (NIH National Cancer Institute). UAS-h*SMPD1* were gifts from Hugo Bellen (Baylor College of Medicine) and Perlara plc respectively.

Heterozygosity of *Smpd1* mutants was determined by the presence of CyO-GFP, which was visualised under the fluorescent dissecting microscope.

### Embryo injections

Embryos were collected at 0-30 minutes of age, manually dechorionated and mounted in heptane glue under halocarbon oil (VWR). Desipramine, myriocin or water was backfilled into 3.5” Drummond capillary tubes and injected with a Picospritzer II microcellular injection unit at a consistent dose using an eyepiece graticule. Embryos were then allowed to develop at 25°C and viable embryos were scored for a tracheal phenotype.

### Fluorescent microscopy

Live embryos were manually dechorionated, selected for genotype, staged on heptane glue on a microscope slide, immersed in halocarbon oil and placed beneath a coverslip. Tracheal sections were then immediately imaged at 1µm z-slices with a 20x or 63x oil objective on a Zeiss 700 or Zeiss 880 confocal microscope.

Lysotracker and DHE staining was conducted as previously described (Evans et al., 2013). In brief, embryos were manually dechorionated, selected for genotype and transferred to Schneider’s insect medium containing 50nM Lysotracker™ Red DND-99 (Thermo Fisher) or DHE (Thermo Fisher), incubated shaking at 700rpm for 30 minutes before mounting in PBS and immediately imaging at 63x on a Zeiss 700 confocal microscope.

### Immunofluorescence

Stage 15/16 embryos were used for immunostaining as the cuticle remained partially penetrant for immunostain uptake at this stage of development. This was performed using a protocol adapted from (Sarkissian et al., 2014). In brief, embryos were dechorionated in 50% Bleach (Acros Organics), washed in phosphate buffered saline-containing 0.1% Tween-20 (PBS-T), and fixed nutating in scintillation vials in 4% paraformaldehyde in PBS-T for 20 minutes beneath a 1:1 Heptane layer. Paraformaldehyde and heptane layers were then removed and replaced by a 1:1 heptane: methanol mix and shaken vigorously for 1 minute. Devitellinised embryos were then washed 3x in methanol, followed by graded washes into PBS-T. Blocking was then performed for 1hr in 5 % horse serum (Gibco) in PBS-T. Embryos were then incubated for 2 days at 4°C in primary antibody, followed by 1hr of PBS-T washes and a further overnight incubation at 4°C in secondary antibody. Embryos were then mounted beneath a coverslip in VECTASHIELD mounting medium with DAPI and imaged on a Zeiss 700 confocal microscope. DSHB 2A12 GASP was the primary antibody (1:200) and the secondary antibody was A10001 goat anti-mouse 488 (1:250).

### Western blotting

3 biological repeats of 50 embryos were collected and flash frozen in liquid nitrogen. Embryos were manually homogenized with a micropestle in NuPage LDS sample buffer and 50nM DTT, run on a NuPage 4-12% Bis-Tris gel (Invitrogen) and transferred to a PVDF membrane. Membranes were blocked in 5% milk powder in TBS-T (TBS with 0.1% Tween-20) for 1 hr and probed by primary antibodies: Lamp1 (Abcam 30687, 1:2000), GABARAP (Abcam 109364, 1:2000), ASP175 (Cell Signalling 9661L, 1:2000) and Actin (Abcam 8224,1:10000) and secondary antibodies goat anti-mouse-HRP (Abcam 6789, 1:10000) or goat anti-rabbit-HRP (Abcam 67211:10000). Membranes were visualised with immobilon crescendo HRP reagent on ImageQuant LAS4000. Bands were normalised to actin and quantified using ImageJ.

## Results

### *CG3376* is the fly ortholog of *SMPD1* required for early development

We first sought to develop a Type A and B NPD model in *Drosophila*, identifying an *SMPD1* ortholog termed CG3376 (Figure 1A). A NCBI conserved domain search identified conserved saposin-like domains and a conserved ASMase domain with 73% sequence homology across the domain (Figure 1B). We henceforth refer to CG3376 as *Smpd1*.

**Figure 1.**
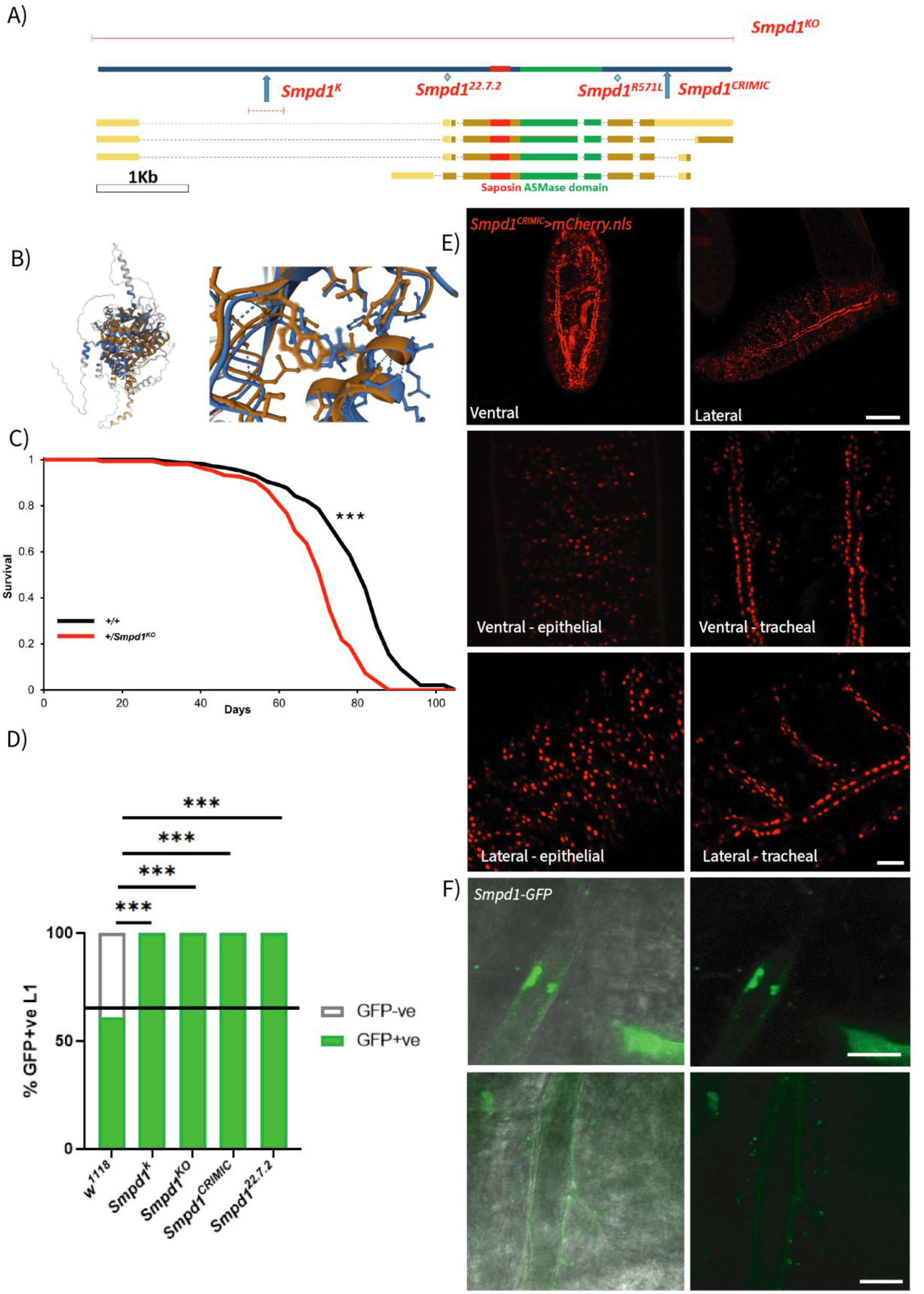
(A) Annotated schematic of the CG3376 coding region and the site of available mutations. (B) Alphafold prediction of human ASM (blue) and the product of *Drosophila* Smpd1 (orange) secondary structures, full view, and a close-up of the ASMase active site. (C) Survival curves of *Smpd1*^*KO*^ heterozygote flies (n=150, Log rank tests p<0.0001). (D) Schematic of crossing scheme and quantification of GFP+ve heterozygous to GFP-ve homozygous 1^st^ instar larvae (n= 272, 293, 151, 403, 113. *w*^*1118*^ vs *Smpd1*^*k/KO/CRIMIC/22*.*7*.*2*^, p<0.0001*** χ^2^ test). (E) Dorsal and ventral views of the localisation of *Smpd1* expression in trachea and epidermis of stage 17 embryos using *Smpd1*^*CRIMIC*^*>mCherry*.*nls*. Scale bar = 100µm for whole embryos, and 25µm for 63x magnification images. (F) Localisation of Smpd*1*-GFP within the tracheal lumen at stage 16 and in vesicles within the tracheal epithelia and on the lumen edge at stage 17, scale bar = 10µm.

A range of CG337*6* mutants were available: an un-excised p-element in the 5’ (*Smpd1*^*k*^), a CRISPR-derived frameshift indel in the 5’ coding region (*Smpd1*^*22*.*7*.*2*^), a full excision of the gene (*Smpd1*^*KO*^) and a truncation mutant caused by insertion of a Gal4-containing transgenic element in the 3^rd^ intron (*Smpd1*^*CRIMIC*^)(Figure 1A). We identified a significant lifespan shortening effect in *Smpd1*^*KO*^ heterozygotes, suggesting a requirement for Smpd1 in *Drosophila* in adulthood as well as development (Figure 1C). All mutants tested were homozygous lethal.

Balancing *Smpd1* mutants over CyO-GFP allowed us to track homozygous flies throughout early development and identify lethal timepoints from which we could infer a requirement for Smpd1 activity. At late stage 17, the developmental stage immediately prior to hatching, we identified morphologically normal *Smpd1*^*-/-*^ embryos exhibiting sequential muscle contractions. However, no viable 1^st^ instar larvae of any *Smpd1* homozygous mutant strain emerged post-hatching (Figure 1D), indicating that loss of Smpd1 activity is lethal at late embryonic stages. As all tested *Smpd1* mutants phenocopy one another, we ascertained that they are all LOF lines and elected to do most of the subsequent experiments with the *Smpd1*^*k*^ mutant.

To identify the tissues requiring Smpd1 function, we tested the expression pattern of Smpd1 at late embryonic development by crossing the *Smpd1*^*CRIMIC*^ line, which contains a Gal4 insertion in the coding region (Lee et al., 2018), to a nuclear localised mCherry. We identified strong ubiquitous staining in both the epidermis and the tracheal luminal cells at stage 17 (Figure 1E). This tracheal expression is corroborated by *in-situ* hybridisations from the Berkeley *Drosophila* Genome Project (BDGP) that reveal that the *Smpd1* transcript in late-stage embryos is enriched in expression in the trachea (Tomancak et al., 2002).

We then studied expression patterns of a GFP-tagged *Smpd1* construct expressed under an endogenous promoter from the fTRG library (Sarov et al., 2016). This identified intraluminal GFP in vacuole-like structures prior to tracheal gas-filling, and then by late stage 17, within vesicles of the tracheal cells (Figure 1F).

### The ASMase activity of Smpd1 is required for tracheal gas-filling

At stage 17, in the hours prior to eclosion, the embryonic trachea in *Drosophila* are morphologically mature, but undergo a transition whereby the liquid layer of the lumen is replaced with a functional gas layer in a sequential cell-specific manner, to allow oxygen exchange and physiological function (Hayashi & Kondo, 2018). As *Smpd1* has strong expression within the trachea, we assessed *Smpd1*^*k*^ mutants under the light microscope to determine if any structural changes in the tracheal system were apparent, preceding the embryonic lethality. This demonstrated that there were severe gas-filling defects in all embryonic lethal *Smpd1* mutants (Figure 2A).

**Figure 2.**
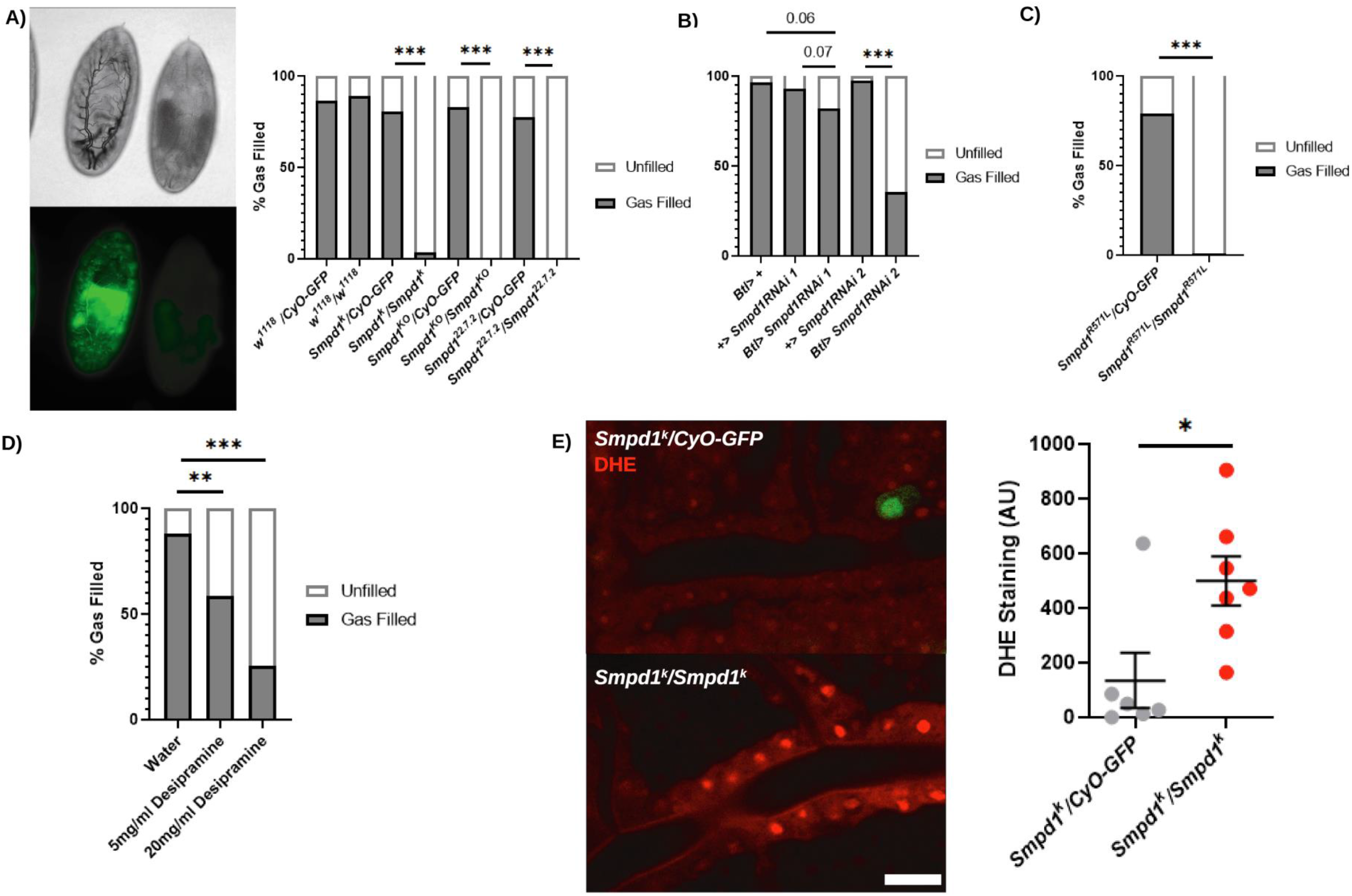
Example brightfield image of stage 17 heterozygous (GFP+ve) and homozygous (GFP-ve) *Smpd1*^*k*^ embryos, demonstrating a loss of gas-filling. (A’) Quantification of gas-filling in *Smpd1* mutants and controls (n=18, 52, 57, 173, 101, 204, 31, 44, *w*^*1118*^ vs *Smpd1*^*k/KO/CRIMIC/22*.*7*.*2*^, p<0.0001, χ^2^ test). (B) Quantification of gas-filling defects following tracheal specific expression (Btl-GAL4) of RNAi against *Smpd1* (n=29, 58, 56, 78, 76, *Smpd1*RNAi 1 vs Btl>*Smpd1*RNAi 1 p=0.07, *Smpd1*RNAi 2 vs Btl>*Smpd1*RNAi 2 p<0.0001, χ^2^ test). (C) Gas-filling defects in *Smpd1*^*R571L*^ embryos containing a point mutation in the ASMase domain (n=220, 95, p<0.0001, χ^2^ test). (D) Quantification of gas-filling defects in stage 17 embryos injected with either desipramine (Des) or water 30 minutes after egg-laying (n=50, 41, 67, PBS vs 5mg/ml Des p=0.0013, PBS vs 20mg/ml Des p<0.0001, χ^2^ test). (E) Example image and quantification of DHE staining in late stage 17 ctrl and *Smpd1*^*k*^ mutant tracheal lumens showing a significant increase in ROS (p=0.0207, t-test), scale bar =10µm.

To confirm this defect was related to tracheal expression of *Smpd1*, we expressed RNAi against *Smpd1* under the control of the tracheal-specific driver Btl-GAL4. This resulted in a mild but statistically significant decrease in gas-filling, suggesting that the loss of tracheal *Smpd1* underlies the gas-filling defect (Figure 2B).

To confirm that this gas-filling defect was dependent on loss of ASMase domain activity, we tested a mutant possessing a CRISPR-generated point mutation (R571L) in the active site of the ASMase domain of *Drosophila Smpd1*, corresponding to the R496L point mutation in human *SMPD1*. This common missense mutation results in a near-complete loss of ASMase activity and is associated with Type A NPD (Levran et al., 1991, Jones et al., 2008). *Smpd1*^*R571*L^ homozygotes robustly showed a loss of tracheal gas-filling (Figure 2C). To further establish that loss of the ASMase domain function is causative for the air-filling defects, we injected desipramine, a potent ASMase inhibitor into healthy blastoderm-stage embryos. This resulted in a statistically significant dose-dependent reduction in tracheal gas-filling at late-stage 17 (Figure 2D). Staining with DHE, a marker of superoxide species production, suggested significantly higher ROS levels in *Smpd1*^*k*^ mutant trachea compared to controls, potentially as a consequence of hypoxia, or an indication of redox defects that may regulate gas-filling (Figure 2E).

### Tracheal endocytosis and luminal clearance are disrupted in *Smpd1*^*-/-*^ embryos

To assess if loss of *Smpd1* causes gross structural tracheal defects, we probed tracheal morphology at stage 15 by staining with an antibody against GASP. This is a secreted chitin-binding protein present in the tracheal lumen, and thus immunostaining for this intrinsic component allows visualisation of the entire tracheal network. Patterns of tracheal branching in stage 15 embryos appeared unchanged between control and mutant flies, with a full complement of tracheal branches and morphologically normal dorsal, lateral, visceral and ganglionic branches (Figure 3A). Previously characterised tracheal defects such as tube overelongation or defects in tube fusion were not observed, suggesting defects in early morphogenesis do not underlie gas-filling defects (Beitel and Krasnow, 2000).

**Figure 3.**
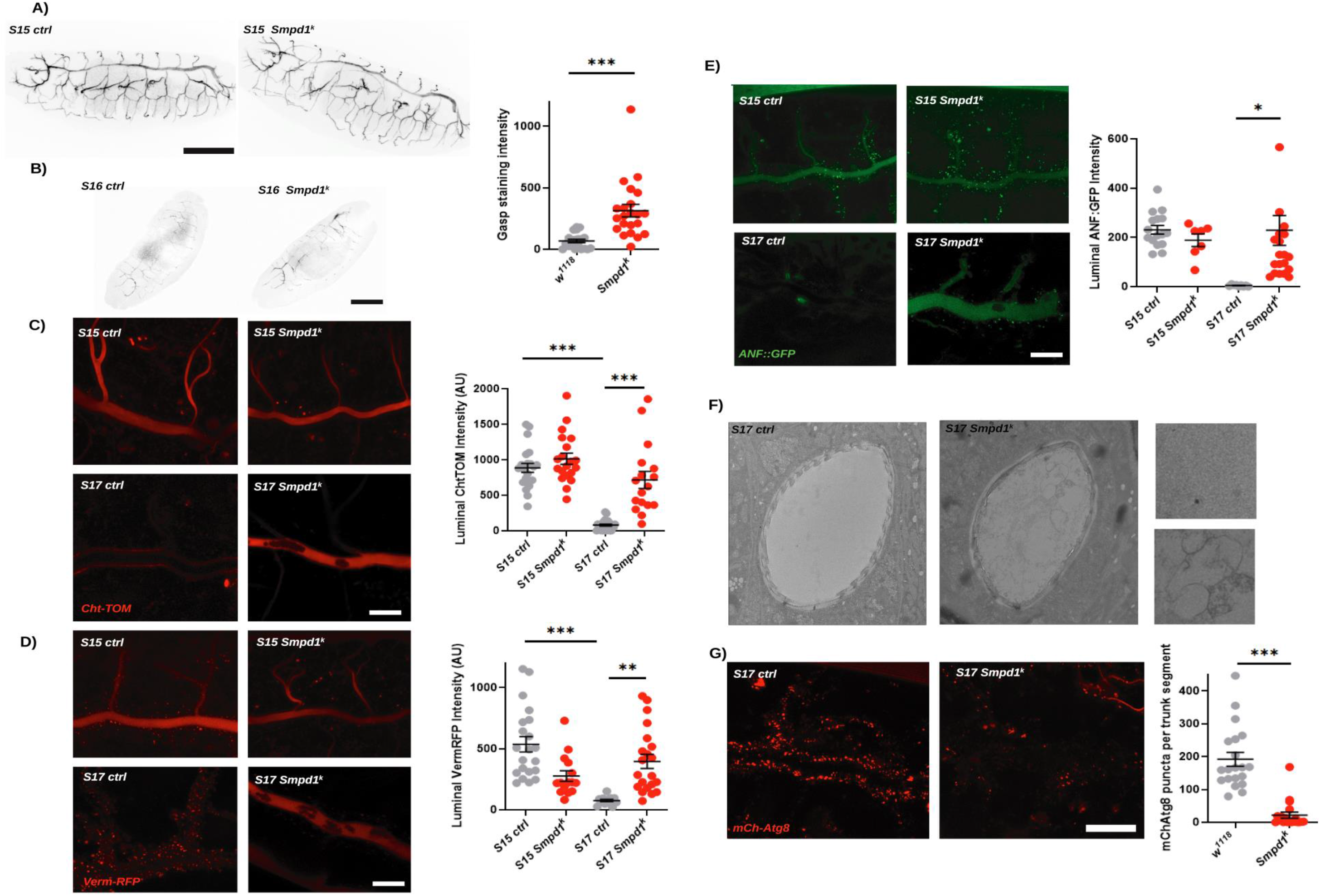
(A) Images of whole tracheal morphology of stage 15 ctrl and *Smpd1*^*k*^ embryos stained for 2A12 (GASP), highlighting that each element of the tracheal network (DT, LT, VT, GT) is intact, scale bar =100µm. (B) Example confocal images of early stage 16 ctrl or *Smpd1*^*k*^ embryos stained for GASP and quantification of luminal staining intensity (p=0.0003, t-test), scale bar = 100µm. (C) Example confocal images of ctrl and *Smpd1*^*k*^ embryos at stage 15 and late stage 17 expressing Btl>VermRFP and quantification of staining intensity across the tracheal lumen (S15 ctrl vs S17 ctrl p<0.0001, S17 ctrl vs S17 *Smpd1*^*k*^ p=0.0015, One-way ANOVA with Tukey’s multiple comparisons test), scale bar = 25µm. (D) Example confocal images of ctrl and *Smpd1*^*k*^ embryos at stage 15 and 17 expressing Btl>ChtTOM and quantification of staining intensity across the tracheal lumen (S15 ctrl vs S17 ctrl p<0.0001, S17 ctrl vs S17 *smpd1*^*k*^ p<0.0001, One-way ANOVA with Tukey’s multiple comparisons test), scale bar = 25µm. (E) Example confocal images of ctrl and *Smpd1*^*k*^ embryos at stage 15 and 17 expressing Btl>ANF:GFP and quantification of staining intensity across the tracheal lumen (S15 ctrl vs S17 ctrl p=0.0233, S17 ctrl vs S17 *Smpd1*^*k*^ p=0.0176, One-way ANOVA with Tukey’s multiple comparisons test), scale bar = 25µm. (F) Transverse TEM section of the dorsal trunk trachea in ctrl or *Smpd1*^*k*^ stage 17 embryos. (G) Example confocal images of ctrl and *Smpd1*^*k*^ embryos at stage 15 and 17 expressing Btl>mCh-ATG8 and quantification of staining intensity (p<0.0001, t-test), scale bar = 25µm.

Tracheal luminal development has been extensively characterised in a temporal fashion. Exocytic events from tracheal luminal cells stimulate development of a chitin exoskeleton, which regulates diameter expansion. As morphogenetic changes are completed, this exoskeleton is then degraded by a pool of secreted chitinases and the resulting milieu of luminal constituents is taken up by tracheal epithelia through endocytosis, presumably resulting in degradation via the endo-lysosomal network (Tsarouhas et al.,2007).

Immunostaining for GASP was increased in *Smpd1*^*k*^ embryos at stage 16, a developmental stage when chitin levels are reduced, indicative of impaired chitin clearance (Figure 3B). We next expressed a fluorescently tagged chitin-binding protein, Cht-TOM, in the trachea of *Smpd1*^*k*^ mutants. In control flies, dispersed chitin staining is visible throughout the tracheal lumen at stage 15, which transitions to the luminal border by late stage 17, as previously described (Tsarouhas et al.,2007). In *Smpd1*^*k*^, however, Cht-TOM remains luminal into late stage 17, with significantly increased fluorescence intensity within the tracheal lumen, indicative of decreased chitin clearance (Figure 3C). We next expressed the chitin-modifying protein, Verm-RFP, which in controls showed a shift throughout development from primarily luminal at stage 15, to a location within intracellular vesicles of tracheal epithelial cells by late stage 17. In *Smpd1*^*k*^, significantly higher expression was retained in the lumen (Figure 3D). These data demonstrate that chitin removal, but not deposition, is disrupted in *Smpd1*^*k*^ mutants.

It has previously been established that clearance of luminal contents, principally degraded chitin and its associated modifiers, is contingent on an endocytic burst (Tsarouhas et al., 2007). To measure this we expressed the secreted factor ANF::GFP in tracheal cells. In controls, ANF::GFP was secreted into the lumen and visible at stage 15, and subsequently cleared by late stage 17, demonstrating both functional exocytic secretion and endocytic uptake. In *Smpd1* mutants, by contrast, ANF::GFP is deposited in the lumen at stage 15 without any visible intracellular aggregates, showing that exocytosis is functional. However, by late stage 17, statistically significant elevation in ANF::GFP fluorescence was observed in *Smpd1*^*k*^ mutant lumens, demonstrating a failure to specifically endocytose luminal constituents (Figure 3E). In late stage 17 *Smpd1*^*k*^ mutants spherical vacuolated structures were visible in the lumen that excluded ANF, Cht or Verm markers, and which were not observed in controls. It is unknown if these structures are membranous or if they originate as secreted vesicles.

To study the ultrastructure of *Smpd1*^*k*^ mutant tracheal lumens in greater detail, we performed transmission electron microscopy (TEM) on transverse sections of stage 17 *Smpd1*^*k*^ mutant and control embryos. In keeping with our fluorescent imaging data, we observed incomplete chitin remodelling at the luminal periphery and a failure to produce distinct taenidial folds at the luminal edge (Figure 3F). In addition to the vacuolated structures seen within the tracheal lumen on fluorescent imaging, there was also evidence of luminal debris. From this data we can conclude that *Smpd1* is required for tracheal development in late embryogenesis, specifically for the clearance of luminal constituents prior to gas-filling.

To further determine the nature of this remodelling defect, and contingent on the findings from ANF::GFP expression experiments that defects were present in tracheal cell endocytosis, we expressed and tracked expression of a range of vesicular markers related to the endo-lysosomal system in the tracheal tissue. Deeper into the endosomal network, expression of the autophagosome marker mcherry::Atg8 revealed a significant decrease in the size and intensity of autophagosomes in *Smpd1*^*k*^ mutant tracheal cells (Figure 3G). This indicates that a disruption of the broader vesicular network may underpin defects in luminal clearance.

### *Smpd1* loss-of-function is associated with changes in ceramide saturation in the absence of abnormalities in lysosomal morphology

As NPD is a lysosomal storage disorder, characterised by lysosomal dysfunction and expansion of the lysosomal compartment (Gabande-Rodriguez et al., 2014), and because we identified defects at other points in the vesicular network, we stained stage 17 *Smpd1*^*k*^ embryos with Lysotracker, a dye specific to acidified vesicles. Curiously, no significant difference in abundance of lysosomal puncta was observed in *Smpd1* mutant tracheal luminal cells compared with controls (Figure 4A). This hints that tracheal defects in *Smpd1*^*-/-*^ embryos do not stem from lysosomal dysfunction, but rather a mechanism of action beyond the lysosome. As Smpd1-GFP appears to be punctate and secreted into the lumen, it is likely it may be secreted and function at the extracellular surface of the plasma membrane.

**Figure 4.**
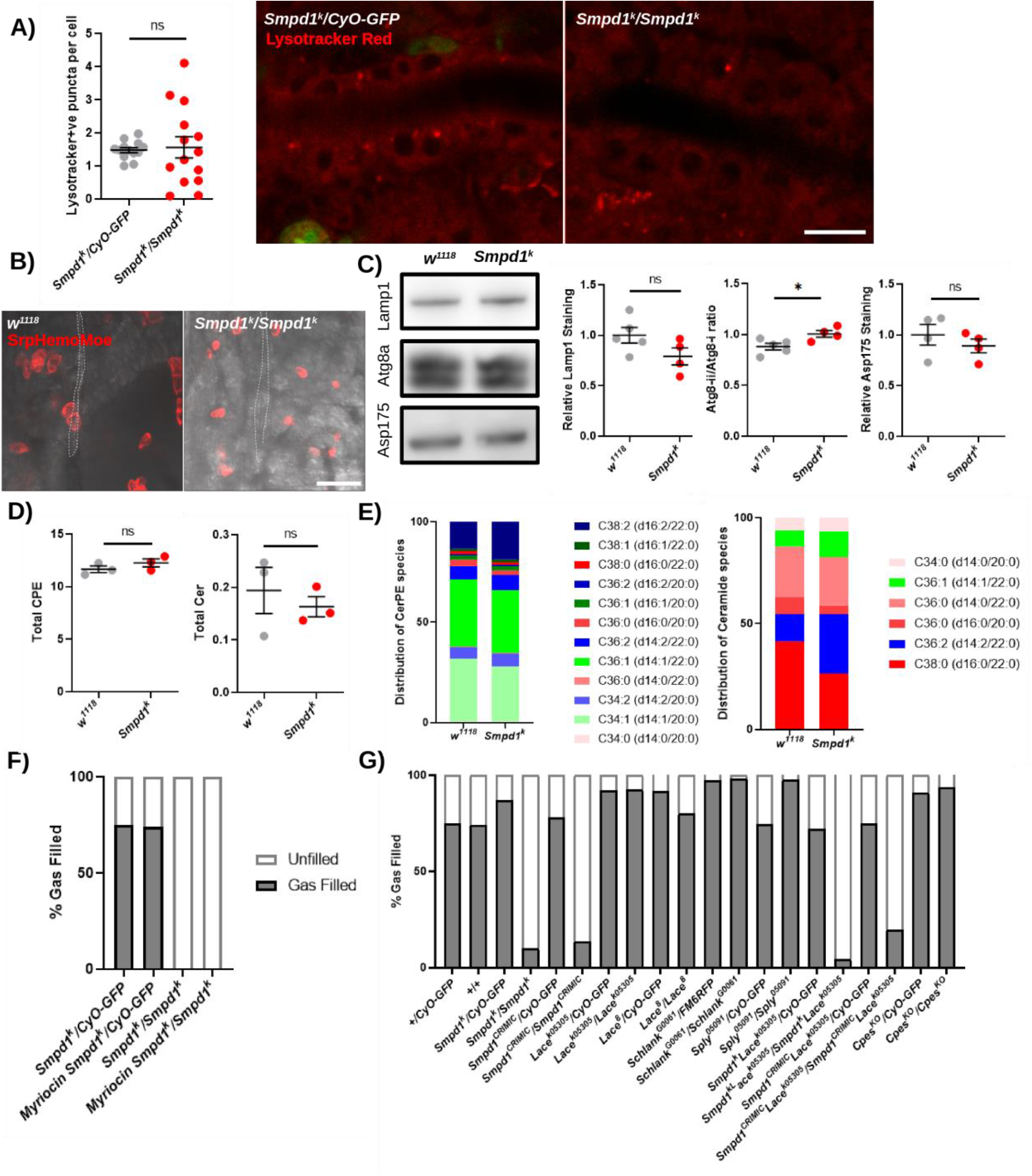
(A) Example images showing Lysotracker Red DND99 staining within S17 ctrl and *Smpd1*^*k*^ tracheal cells and quantification of (p=0.8113, t-test), scale bar=10µm. (B) Representative images of macrophages (Srp.Hemo.Moe.mCherry) and trachea (Brightfield) showing no colocalisation or infiltration in ctrl or *Smpd1*^*k*^, scale bar =25µm. (C) Western blots on whole stage 17 embryo extracts for ctrl or *Smpd1*^*k*^. Example images and quantification normalised to actin for (C’) Lamp1 (p=0.1094. t-test), (C’’) Atg8a-II/I ratio (p=0.003, t-test) and (C’’’) Asp175 (p=0.4025, t-test), demonstrating no change in embryo-wide lysosomal or apoptotic activity. (D) Graphs of overall abundance of Ceramide (Cer) and Cer phosphoethanolamine in ctrl or *Smpd1*^*k*^ stage 17 embryos (ctrl vs *Smpd1*^*k*^ CPE p=0.2943, ctrl vs *Smpd1*^*k*^ Cer p=0.5532, t-test) as detected by HPLC. (E) Graph of sphingolipid species by chain length and saturation state in ctrl or *Smpd1*^*k*^ stage 17 embryos. (F) Quantification of gas-filling defects in *Smpd1* flies with myriocin or DMSO supplementation to maternal diet. (G) Quantification of gas-filling defects in *Smpd1* or mutations in sphingolipid biosynthesis.

To assess if there was any macrophage involvement in respiratory pathologies, as is the case in NPD, we visualised hemocytes with the cytoskeletal marker Hemo-Moe.mCherry and failed to find any instance of macrophages contacting or infiltrating the trachea in either control or *Smpd1* mutants (Figure 4B).

Furthermore, we conducted western blots on whole embryos at late stage 17 and did not identify any change in the levels of a common lysosomal marker, LAMP1 (Figure 4C). This supports the fact that bulk lysosomal defects are not present in the tracheal tissue of *Smpd1* deficient mutants. Similarly, there was no significant difference in staining for active Caspase 3, a marker of apoptosis, demonstrating that widespread cell death does not occur in *Smpd1*^*k*^ mutants (Figure 4C). A significant increase was detected in both Atg8aII and Atg8aII/Atg8aI ratio, which is indicative of increased autophagosome formation, likely related to hypoxic tissue degradation (Figure 4C).

*Drosophila* lack the Smpd1 substrate SM, possessing instead the structural analog CPE. We assessed abundances of CPE and its putative derivative, ceramide (Cer), using targeted mass spectroscopic quantitative lipidomic analysis (Kunduri et al., 2018). The overall abundance of Cer or CPE was not altered in *Smpd1* embryos compared to controls (Figure 4D). Similarly, no significant differences in chain length or saturation were evident in classes of CPE, suggesting that CPE is unchanged by the loss of Smpd1 function. Significant changes were, however, identifiable with Cer species, with an increase in unsaturated Cer in *Smpd1*^*k*^ mutants (Figure 4E). The mechanism of this shift and its functional relevance is unknown, but unsaturated sphingolipids have decreased density and interaction with cholesterol within a membrane, which may indicate a change in lipid raft stability at the membrane and consequent effects on vesicle dynamics (Pinto et al., 2011, Maula et al., 2015).

This data suggests that the nature of the defects observed in *Smpd1*^*-/-*^ mutants departs from canonical NPD pathology, with no obvious change in lysosomal area or sphingolipid abundance. It is feasible that minor effects of *Smpd1* deficiency in the trachea are not detectable in a whole fly homogenate. However, more likely is the possibility that Smpd1 is secreted, as has been proposed elsewhere (Tsarouhas et al., 2023). Smpd1 may act on CPE at the membrane to facilitate signalling events at a level imperceptible to bulk biochemical analysis.

Lysosomal lipid salvage pathways are thought to be a major source of cellular Cer, and it is thought that phenotypes may therefore result from a local depletion of Cer, in addition to an overabundance of CPE (Schuchman, 2010). To determine if the gas-filling defects are related to Cer abundance, we raised maternal flies on fly media containing myriocin, a potent serine-palmitoyltransferase (SPT) inhibitor that blocks de-novo ceramide synthesis. This manipulation had no effect on the gas filling defects of *Smpd1*^*k*^ embryos (Figure 4E).

To directly measure the impact of zygotic sphingolipid synthesis on gas-filling, we screened and found no gas-filling defects in embryos mutant for *Lace (SPT), Schlank* (ceramide synthase*), Sply* (sphingosine phosphate-1-lyase) and *CPES* (ceramide phosphoethanolamine synthase), required for the biosynthesis of serine-palmitoyltransferase, ceramide, sphingosine-1-phosphate and CPE respectively (Figure 4F). Knockdown of SPT using the *Lace*^*k05305*^ mutant in *Smpd1*^*k*^ mutants, resulting in a condition of presumptively lower Cer, did not alter the gas-filling defects and had no effect on control gas-filling (Figure 4F).

### The gas filling defects seen in *Smpd1* deficient flies are rescued by a combination of restricting dietary lipids and zygotic ceramide biosynthesis

Our data suggests *Smpd1* is required in an extra-lysosomal manner for clearance of tracheal proteins, and as an ostensible lipid hydrolase. We therefore hypothesised that *Smpd1* might modify lipid constitution at the membrane. We thus reasoned that modulating the levels of certain lipids in plasma membranes might influence the *Smpd1* gas-filling defect.

As our targeted lipidomics failed to compellingly point to a particular lipid substrate for Smpd1, we aimed first to disrupt bulk lipid bioavailability. As maternal synthesis, zygotic synthesis and dietary presence of lipids could all influence lipid availability, we began a combinatorial approach to shift membrane lipid constitution. We made the following assumptions about lipid homeostasis in the late embryo: A) zygotic lipid constitution will be influenced by maternal deposition of lipids; and B) egg production occurs in adulthood so the deposition of lipids in the ovaries will be influenced by adult maternal diet and lipid synthesis. We raised *Smpd1*^*k/+*^ heterozygote mothers on a holidic pre-defined diet to selectively add and remove elements from the diet (Piper et al., 2014). As *Drosophila* are sterol auxotrophs (Clayton, 1964, Carvalho et al., 2010), the only lipid in holidic medium is cholesterol, and all other lipids in the embryo have to be de-novo synthesised by maternal or zygotic synthesis pathways. To our surprise, raising maternal *Smpd1*^*k*^ heterozygote flies on a holidic medium for 6 days was sufficient to uncover a robust population of *Smpd1*^*k*^ homozygote mutant progeny with gas-filled trachea (Figure 5A).

**Figure 5.**
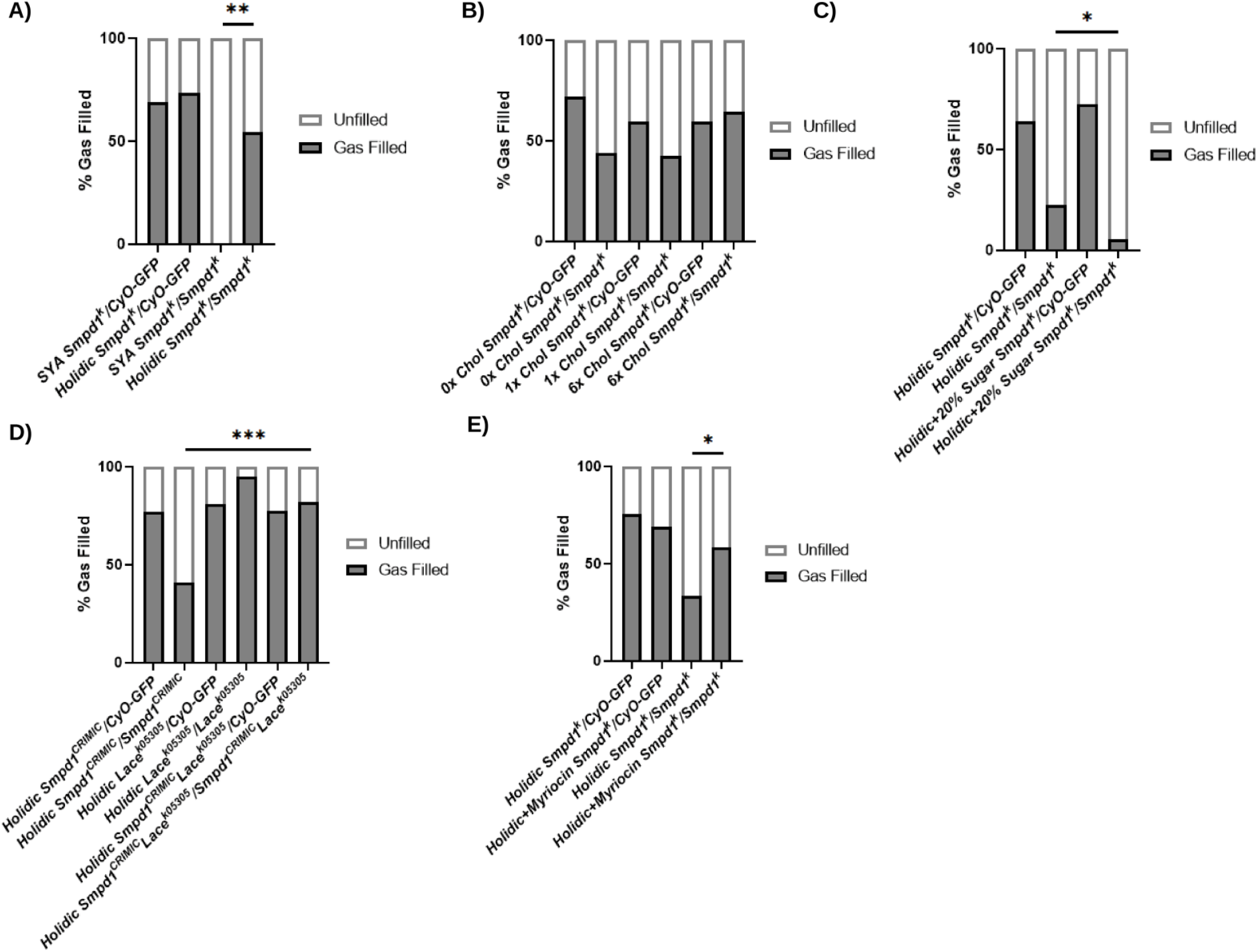
(A) Quantification of gas-filling defects in ctrl or *Smpd1*^*k*^ Stage 17 embryos from flies raised on an SYA or holidic diet (n= 45, 83, 16, 33, SYA *Smpd1*^*k*^ vs holidic *smpd1*^*k*^ p=0.0002, χ ^*2*^ test). (B) Quantification of gas-filling defects in ctrl or *Smpd1*^*k*^ Stage 17 embryos from flies raised on an holidic diet with 0x, 1x or 6x dietary cholesterol, showing no significant effect (n=43, 64, 67, 16, 26, 28, 0x Chol *Smpd1*^*k*^ vs 1x Chol *Smpd1*^*k*^ p=0.9269, 1x Chol *Smpd1*^*k*^ vs 6x Chol *Smpd1*^*k*^ p=0.1056, χ ^*2*^ test). (C) Quantification of gas-filling defects in ctrl or *Smpd1*^*k*^ Stage 17 embryos from flies raised on an holidic diet with 0 or 20 % sugar (n= 106, 53, 91, 36, p=0.0298, χ ^*2*^ test). (D) Quantification of gas-filling defects in ctrl, *Smpd1*^*k*^ Stage 17 embryos from flies combining *Lace*^*K05305*^ raised on an holidic diet (n= 99, 34, 120, 41, 195, 78, *Smpd1*^*CRIMIC*^ vs *Smpd1*^*CRIMIC*^ *Lace*^*K05305*^ p<0.0001, χ ^*2*^ test). (E) Quantification of gas-filling defects in ctrl or *Smpd1*^*k*^ stage 17 embryos from maternal flies raised on an holidic diet with or without myriocin supplementation in the maternal diet (n=199, 156, 86, 77, *Smpd1*^*k*^ ctrl vs *Smpd1*^*k*^ myriocin p=0.0016, χ ^*2*^ test).

As sterols are a key constituent of lipid rafts and may form microdomains with sphingolipids at the membrane, we removed or over-supplemented cholesterol in holidic diets at 0x, 1x and 6x cholesterol, and found no significant difference in gas-filling in *Smpd1*^*k*^ mutants with changes in sterol supplementation (Figure 5B). To determine if the gas-filling rescue was related to lipid bioavailability, we supplemented holidic medium with 20% sugar, which is a substrate for lipid synthesis following glycolysis and fatty acid synthesis. This showed that sugar supplementation significantly abrogated gas-filling in *Smpd1*^*k*^ on holidic media (Figure 5C)(Musselman et al., 2011). Direct supplementation of the holidic medium with 3% coconut oil or lard resulted in a complete cessation of egg-laying. Thus, we conclude from these findings that maternal dietary lipids are required for the maintenance of lipid membranes and that *Smpd1* is an essential regulator of membrane remodelling and homeostasis.

To assess if gas-filling defects were related to overabundance of Cer species, we inhibited zygotic Cer synthesis using a *Lace*^*k05305*^; *Smpd1*^*k*^ double mutant. Whilst loss-of-function *Lace* mutations by themselves had no deleterious effect on gas-filling, in combination with a holidic medium, *Lace*^*k05305*^; *Smpd1*^*k*^ double mutants had significantly higher gas-filling percentage than *Smpd1*^*k*^ mutants. These findings suggest that lowering Cer worsens the tracheal defects in *Smpd1* mutant embryos (Figure 5D). To further test this hypothesis, we incorporated myriocin into the maternal holidic diet and found that this significantly increased the gas-filling rescue of the holidic diet in *Smpd1* mutants (Figure 5E). Taken together, this data suggests that an overabundance of a Cer derivative leads to a gas-filling tracheal defect, which requires Smpd*1* ASMase activity to overcome. Thus, Cer lowering offers a potential therapeutic strategy for the treatment of Type A and B NPD and associated disorders.

## Discussion

Type A and B NPD is a rare disorder with few effective treatment options. Here we present a novel NPD fly model lacking the *Drosophila* orthologue of *Smpd1*.

Similar to patient reports, we demonstrated that *Smpd1* mutants have defects in tracheal luminal clearance. As macrophages do not infiltrate the trachea in *Drosophila* and were not seen to do so in *Smpd1* mutants, it is likely that tracheal cells, known to be endocytically active, have an analogous function to macrophages in mammals in turning over luminal constituents. The absence of lysosomal or macrophage phenotypes in tracheal cells raises the possibility that respiratory defects in humans may not arise solely from canonical lysosomal storage defects, but perhaps through alternative pathologies related to secretory functions of ASM.

*Smpd1* mutant embryos lack several of the key hallmarks of NPD, in lysosomal dysfunction and sphingolipid accumulation. However, this model has the potential to uncover previously understudied mechanisms of disease progression. Our data suggests extracellular defects in the tracheal lumen and targeted effects on membrane dynamics are important for developmental phenotypes. Thus, Smpd1 displays a secreted rather than a lysosomal role in *Drosophila* embryos. Evidence of an increase in autophagosomes is in keeping with other *Drosophila* lysosomal storage disorder (LSD) models, underscoring common mechanisms between diverse LSDs (Atilano et al., 2023, Hull et al., 2024). Furthermore, *Smpd1* is abundantly expressed in adulthood in the glia and fat body, suggesting important roles in adulthood, and adult-specific knockout of *Smpd1* may produce more canonical LSD phenotypes.

Whilst pulmonary disorders are commonly reported across NPD types, their aetiology is under-investigated and divergent. One study found uniform pulmonary involvement manifesting either as chronic obstructive pulmonary disorder or chronic respiratory failure (Guillemot et al.,2007), and another showed 90% of Type A NPD patients died of respiratory failure (Mcgovern et al., 2006). In the milder form of the disease, Type B NPD, a high proportion of patients showed evidence of lung dysfunction (Mcgovern et al., 2008). Only individual case studies have specified respiratory pathology, although foam cells within the airway are frequently identified (Guillemot et al.,2007), alongside Interstitial lung disease (Opoka et al., 2020), pulmonary alveolar proteinosis (Sideris and Josephson., 2016), and macrophage infiltration. An ASM knockout mouse model demonstrated a “foam” phenotype macrophage accumulation within the airways, associated with crystalline aggregation of chitinase-like-protein YM1 (Poczobutt et al., 2021).

Our genetic and pharmacological data using a *Lace* mutant and myriocin strongly points towards abundance of Cer or a Cer derivative as being restrictive to gas-filling. It has previously been shown that CPE is an SM analogue in *Drosophila*, required for the formation of lipid rafts and their most abundant constituent (Kunduri et al.,2018). Lipid rafts are key to major membrane-trafficking processes such as clathrin-independent endocytosis, which relies on a specialised type of lipid raft, caveolae, in addition to diverse roles anchoring and organising membrane proteins. ASM has been shown to induce spontaneous vesicle formation in artificial liposomes through the generation of ordered Cer domains (Holopainen et al., 2000). Furthermore, exocytosed ASM is required for injury-dependent endocytosis at the plasma membrane, and ASM-deficient cells are endocytosis-deficient (Tam et al., 2010). We have shown ANF:GFP, a secretion marker, is not cleared from *Smpd1*^*k*^ tracheal lumens, suggesting a defect in endocytic uptake of luminal constituents. It is possible that Smpd1 functions at the tracheal membrane in *Drosophila* in a similar manner to ASM in cell lines, to co-ordinate membrane budding events.

One shortcoming of our work is we have not directly shown *Smpd1* cleaves CPE. Thus, it is conceivable, as CPE levels are not increased, that *Smpd1* may target other sphingolipids. Though previous lipidomic analyses have failed to identify native SM in *Drosophila*, including in embryos, it is possible SM derived from the diet could be present and functional in understudied tissues (Rietveld et al., 1999, Seppo et al., 2000, Guan et al., 2013, Kunduri et al., 2018). As point mutations in the active site of ASMase cause gas-filling defects, it is likely the enzyme functions through cleavage of a SM-like lipid. Intriguingly, the chitin subunit N-acetylglucosamine is a common lipid modification, and membrane-chitin meshworks may be involved in structuring chitin remodelling. *Smpd1* may be developmentally required to cleave chitin-modified sphingolipids in the tracheal lumen.

Another lipid-modifying enzyme that has been implicated in tracheal gas-filling is *Wunen*, a secreted lipid phosphate phosphatase that principally dephosphorylates sphingolipids such as sphingosine-1-phosphate and Cer-1-phosphate. However, *Wunen* loss of function mutants have different tracheal phenotypes to *Smpd1*, showing defects in lumen fusion and accumulation of luminal proteins, suggestive of the fact that they are not regulating the same pathway (Ile et al., 2012).

Whilst this manuscript was in preparation, a similar paper describing tracheal defects in *Smpd1* (*dASM*) was published (Tsarouhas et al., 2023), focusing on mechanisms of surfactant secretion in the lumen. In concordance, we also see vacuolated structures in the lumen of *Smpd1*^*k*^ mutants through TEM and fluorescent imaging. As defects in the number and location of various intracellular vesicles are present in *Smpd1*^*k*^ tracheal cells, we speculate that broad defects in vesicle sorting occur in these cells, likely contingent on a dysregulation of membrane microdomains required for these events.

Crucially, the gas-filling phenotypes we identify are visible by light microscopy and require little preparation; as a consequence, mutants can be quickly screened for genetic, dietary or pharmacological modifiers and has potential as a tool to uncover novel therapies for Type A and B NPD. Using this model, we have demonstrated that lowering Cer levels, via genetic or pharmacological means, when flies were reared under a maternal diet lacking ceramides, led to a reversal of the gas-filling tracheal defects. Thus, ceramide lowering represents a potential therapeutic strategy for Type A and B NPD. Moreover, myriocin, a Cer lowering drug, is already FDA-approved for use in humans. Thus, this work highlights the potential of re-purposed drugs like myriocin to be fast tracked into clinical trials in Type A and B NPD.

## Acknowledgements

We wish to thank Jairaj Acharya for sending cpes^KO^ and Uas-cpes fly lines, Hugo Bellen and Perlara Plc for sending Uas-hSmpd1 and Giulia Mastroianni and the QMUL electron microscopy facility for assistance with TEM. This work was supported by the Wellcome Trust (Wellcome Trust Career Development Fellowship, 214589/Z/18/Z to KJK) and funding from the Rosetrees Trust (M701 and M701A to KJK).

